# Altered kinematics and muscle activities during half squats in people with ankle sprain history

**DOI:** 10.1101/2025.06.10.658971

**Authors:** Umi Matsumura, Toshiya Tsurusaki, Rena Ogusu, Shimpei Yamamoto, Yeonghee Lee, Shinya Sunagawa, Hironobu Koseki

## Abstract

Regardless of the high recurrence rate, copers manage the damage after a sprain. Copers do not experience ankle instability and continue their daily activities. However, it is unclear whether copers can use movement strategies similar to people without an ankle sprain. Identifying movement strategies can improve efficacy and prevent unwanted effects during training. Therefore, we aimed to compare the kinematics and muscular activities of the lower limbs during half squats and gait between copers and healthy controls. Ten copers (7 men, 3 women; age = 21.6 ± 1.1 years) and 10 controls (6 men, 4 women; age = 22.5 ± 2.8 years) were included in the cross-sectional study. We compared angles of the thigh, shank, and foot calculated from inertial measurement units and muscle activities of the biceps femoris long head, rectus femoris, gastrocnemius, and tibialis anterior between copers and controls during half squats and gait. For half squats, copers demonstrated thigh restriction with an effect size of 0.67–1.35, whereas we observed no significant differences in the shank and foot angles in the descending phase. Copers demonstrated less activity with an effect size of 0.61–1.58 in all four muscles. Copers showed lower activities of the rectus femoris and tibialis anterior during the same period of thigh restriction. For gait, there were no significant differences in lower limb angles and muscle activities between copers and controls. Copers may restrict thigh rotation unintentionally, leading to less muscle activity during half squats. Removing unconscious restrictions using sensor-based quantitative feedback can enable copers to obtain more efficient training. Our results suggest that assessing the limb movements of copers is important for efficient training, even if they do not experience instability to continue with their daily activities.

## Introduction

Lateral ankle sprain is a common musculoskeletal injury during sports and daily activities [1]. Following an initial ankle sprain, a high percentage of individuals experience a recurrent sprain [1], and 40% develop chronic ankle instability (CAI) [2]. CAI is characterized by persistently repeated episodes of giving way and recurrent injuries, accompanied by pain, swelling, limited motion, muscle weakness, and decreased self-reported function [3]. Individuals with CAI experience difficulties in performing daily and sports activities [4]. In addition, repetitive cartilage degeneration due to CAI is a potential risk factor for developing osteoarthritis [3].

After the initial ankle sprains, damaged proprioceptors of the sprained ankle affect neuromuscular strategies of the ankle and the proximal joints [3, 5]. Altered neuromuscular strategies in people with CAI have revealed abnormalities in muscle activities indicated by surface electromyography (EMG) during functional tasks [1, 6, 7] and gait [5, 8, 9]. People with CAI have also demonstrated altered kinematics, such as increased plantarflexion or reduced dorsiflexion, decreased knee flexion, and increased hip-flexion angles during gait [9, 10]. These altered kinematics may be the compensatory strategy of the lateral ankle sprain [8]. It may cause movements to be performed less efficiently and contribute to poor dynamic stability, which causes recurrent injuries [9].

Regardless of the high rate of recurrent injuries after the initial ankle sprain, some people do not experience subsequent injuries or giving way episodes. Since these people manage to cope with the damage that could lead to CAI after ankle sprains, they are referred to as "copers” [10]. Copers experience no self-reported ankle instabilities during activities of daily living and sports and resume moderate physical activity [10]. In addition, almost half of the individuals with a history of ankle sprain do not seek medical attention and do not receive proper treatment or rehabilitation [11].

However, compared to controls with no history of ankle sprains, copers have demonstrated altered joint kinematics and muscle activities during jump landing [12] and single-leg standing [13]. It is questionable whether copers can use the same movement strategies as people without a history of ankle sprains, even if they continue their daily activities. Since altered movement strategies can cause limb instability or less movement performance and efficacy [14], it is important to assess the movement strategies of copers to prevent recurrent sprain and continue exercise effectively.

The gold standard for assessing movement strategies is motion analysis using a camera-based three-dimensional motion capture system and force plates in a laboratory [8–10]. Recently, inertial measurement units (IMUs) comprising accelerometers, gyroscopes, and magnetometers have been used to capture limb kinematics because they can overcome some limitations of the clinical use of tests in laboratory settings, such as cost, lack of portability, and requirement of a sensitive measurement environment [15–17]. Wearable sensors such as EMG and IMUs enable quantitative analysis in clinical or sports settings.

The squat is a fundamental exercise in the field of strength and conditioning [18, 19]. Due to the similarity of its biomechanics and muscle activities to a wide range of daily activities and sports movements, squatting exercises are frequently chosen as the therapeutic intervention in multiple fields for improving quality of life, training lower limb muscles [20], and enhancing sports performance [21]. Identifying kinematics and muscle activity changes during squat exercises can improve the training effect and prevent unwanted training effects in the intervention [19–22]. To our knowledge, no study has examined the kinematics and muscle activities of copers during squats using wearable sensors. Therefore, we aimed to compare the kinematics and muscle activities of the lower limbs during squats and gait between copers and healthy controls using wearable sensors. We chose a half squat rather than a full squat to minimize head and trunk movements [19]. We focused on the limb rotational angles, not joint angles, as the kinematics parameters to assess the relationship between movement direction and gravity. We hypothesized that altered limb angles and muscle activities would be observed in copers for half squats and gait.

## Materials and Methods

### Study Design

This was a secondary cross-section analysis conducted in Nagasaki University Graduate School of Biomedical Sciences between October 1st 2018 to October 31st 2022.

### Participants

The participants were ten copers and ten age- and body-shape-matched healthy controls. All participants provided written informed consent after receiving an explanation of the study. We conducted this study per the Helsinki Declaration’s guidelines, and the Ethics Committee of the Graduate School of Medical Facilities, Nagasaki University, Nagasaki, Japan (Approval number 18061429, 20070905, 21090905).

The inclusion criteria for the coper group were 1) a history of at least one significant ankle sprain with inflammatory symptoms that led to a minimum of one day of interrupted physical activity at least 12 months before enrolling in the study; 2) having more than 1.5 h of physical activity per week; 3) no recurrent ankle sprains within the 3 months before the study; and 4) no previous “giving way” and/or “feelings of instability” on the injured ankle. Feelings of instability were based on an ankle instability instrument: a “yes” answer to the question “have you ever sprained an ankle?” and a "no" answer to the questions regarding perceived ankle instability during activities of daily living such as walking or climbing stairs [10]. Inclusion criteria for the control group were 1) no history of an ankle sprain; 2) no history of the ankle giving way; 3) no orthopedic or neurological disorders; and 4) having more than 1.5 h of physical activity per week. Exclusion criteria for both groups were: 1) a history of other musculoskeletal diseases or neurological diseases; 2) pain, inflammation, and swelling that interferes with motions; 3) acute sport-related injury to the lower limb that required rest from physical activities 3 months before the study; and 4) history of surgery or rehabilitation in the lower limb [9].

### Procedures

Participants wore shorts and were barefoot. We instructed the participants to keep their head and trunk in line with the hip joints to minimize the effects of head and trunk movements during the half squat [19]. We fixed the leg width during the squat at shoulder width, and the squat was carried out at a comfortable speed for the participants with the arms folded in front of the chest. Participants walked straight for 5 m in their natural walking form for the gait task at a comfortable speed while looking forward. All participants performed half-squat and gait tasks thrice. A 1-min rest between trials and a 5-min rest between tasks were allowed to minimize fatigue. Following the trial, the participants performed a maximum voluntary isometric contraction (MVIC) task.

The MVIC trials for each muscle were performed based on the specific limb positions used in manual muscle testing [23]. We instructed the participants to exert a maximal contraction and hold it for 3 s while the physical therapist applied resistance and provided verbal encouragement, ensuring maximal effort was exerted. Mean values of 0.5 s during maximum effort for 3 s were used as a reference MVIC value [7].

### Data Acquisition

We examined the sprained legs of the copers and the right legs of the controls. We calculated lower limb rotation angles using IMUs (LP-WSD1101-OA, 5G/ 300dps, LOGICAL PRODUCT, Fukuoka, Japan) sampled at 1000 Hz. We mounted the IMUs on the lateral aspect of the thigh, shank, and foot using double-sided adhesive tape to measure limb rotation in the sagittal plane. The attachment positions were the thigh at halfway between the greater trochanter and lateral femoral condyle, the shank at halfway between the lateral femoral condyle and lateral malleolus, and the middle of the dorsum of the foot [16, 17].

We measured muscle activation of the biceps femoris long head (BFL), rectus femoris (RF), gastrocnemius (GAS), and tibialis anterior (TA) using surface EMG (LP-IW2PAD; LOGICAL PRODUCT, Fukuoka, Japan) sampled at 1000 Hz with a 4-channel system (LP-WSD1002-OA; LOGICAL PRODUCT). We focused on the major muscles that represent ankle and knee movements in the sagittal plane, such as ankle dorsiflexion/plantar flexion and knee extension/flexion. We positioned bipolar Ag/AgCl EMG electrodes (VL-00-S/25; METS, 1-7, Tokyo, Japan) according to the Surface EMG for Non-Invasive Assessment of Muscles guidelines [24] with an interelectrode distance of 30 mm, which was significantly wider than the standard 20 mm, considering contraction errors in the clinical settings [25]. Before attaching the electrodes, we shaved the skin and disinfected it with alcohol, and a physical therapist palpated the muscle valleys to prevent artifacts from muscle crosstalk.

### Data Processing

Data from the IMUs and EMG were time-axis synchronized using a wireless 8-channel logger (LP-WSD1311-OA; LOGICAL PRODUCT) and downsampled to 1/30 s, a video camera scale [26], assuming limb angles can be evaluated from video cameras.

The limb angle in the sagittal plane was calculated by integrating the angular velocity in the sagittal plane measured using the IMUs with time displacement [16]. We applied a Kalman filter to minimize the common integration error in IMUs, through which the signal was low-passed at 6 Hz, using a zero-phase fourth-order Butterworth filter [15, 17].

EMG signals were processed as follows: band-pass filtered at 20–450 Hz, full-wave rectified, low-pass filtered at 6 Hz with a fourth-order zero-lag Butterworth filter, and normalized to the reference MVIC value [19].

For both groups, the mean values of three trials in half squats and gait were used for analysis. We removed the first two steps from the beginning, and one gait cycle from the third step was cut and used for analysis. Data for one squat and one gait cycle were converted to 100%. For the half-squat, the descending phase was defined when the shank angle reached its peak, and the ascending phase was defined after reaching the peak. The descending phase lasted 60% of the movement duration, followed by the ascending phase. For the gait, we detected the beginning of the stance and swing phases according to the methods of Watanabe et al [17]. In brief, if the sum of the absolute value of acceleration signals of the three axes exceeded 0.15, it was considered the beginning of the stance. If it fell below 0.15, it was considered the beginning of the swing phase. The stance phase lasted 60% of the movement duration, followed by the swing phase.

### Statistical Analyses

We calculated the sample size according to Koshino et al. [8] using a significance level of 0.05, a detection power of 80% and effect size of 0.5 by using G*Power 3.1. Eight participants per group were expected to detect a group difference during the task. We included 10 participants per group to compensate for the possibility of defective data. We used JMP Pro 15 software (SAS Institute, Tokyo, Japan) for statistical analysis. We used the Shapiro–Wilk test to confirm the normal data distribution. We used two-tailed independent t-tests to identify group differences in limb angles and muscle activity. Statistical significance was set at *P* < 0.05. In addition, we calculated Cohen’s d effect size (ES) to evaluate the magnitude of group differences. We interpreted ES as follows: ≥0.80, large; 0.50–0.79, moderate; 0.20–0.49, small; and <0.20, trivial [5, 6].

## Results

There were no significant group differences between copers and controls regarding age, height, or body weight (copers: 7 men, 3 women; age 21.6 ± 1.1 years, height 1.68 ± 0.10 m, weight 63.1 ± 7.0 kg and healthy controls: 6 men, 4 women; age 22.5 ± 2.8 years, height 1.67 ± 0.10 m, weight 63.0 ± 15.1 kg). Copers had a previous ankle sprain an average of 4.9 ± 5.0 years ago.

For half squats, copers revealed a significantly decreased thigh rotation angle in the descending phase compared to that in controls *(P* < 0.001) with moderate to large ES (d = 0.67–1.35) (Fig 1). The mean group difference was 10.3°. Copers also demonstrated significantly lower foot rotation angles from 70% of half squats *(P* < 0.001) than controls with moderate ES (d = 0.63–0.72) (Fig 1). The mean group difference was 2.2°. There were no significant differences *(P* > 0.05) in the shank angle (Fig 1). Copers demonstrated significantly less mean activity in the descending phase *(P* = 0.001-0.35) with moderate to large ES (d = BFL: 0.83–1.40, RF: 0.66–1.58, GAS: 0.7, TA: 0.62–1.30) (Fig 2).

**Fig 1.**
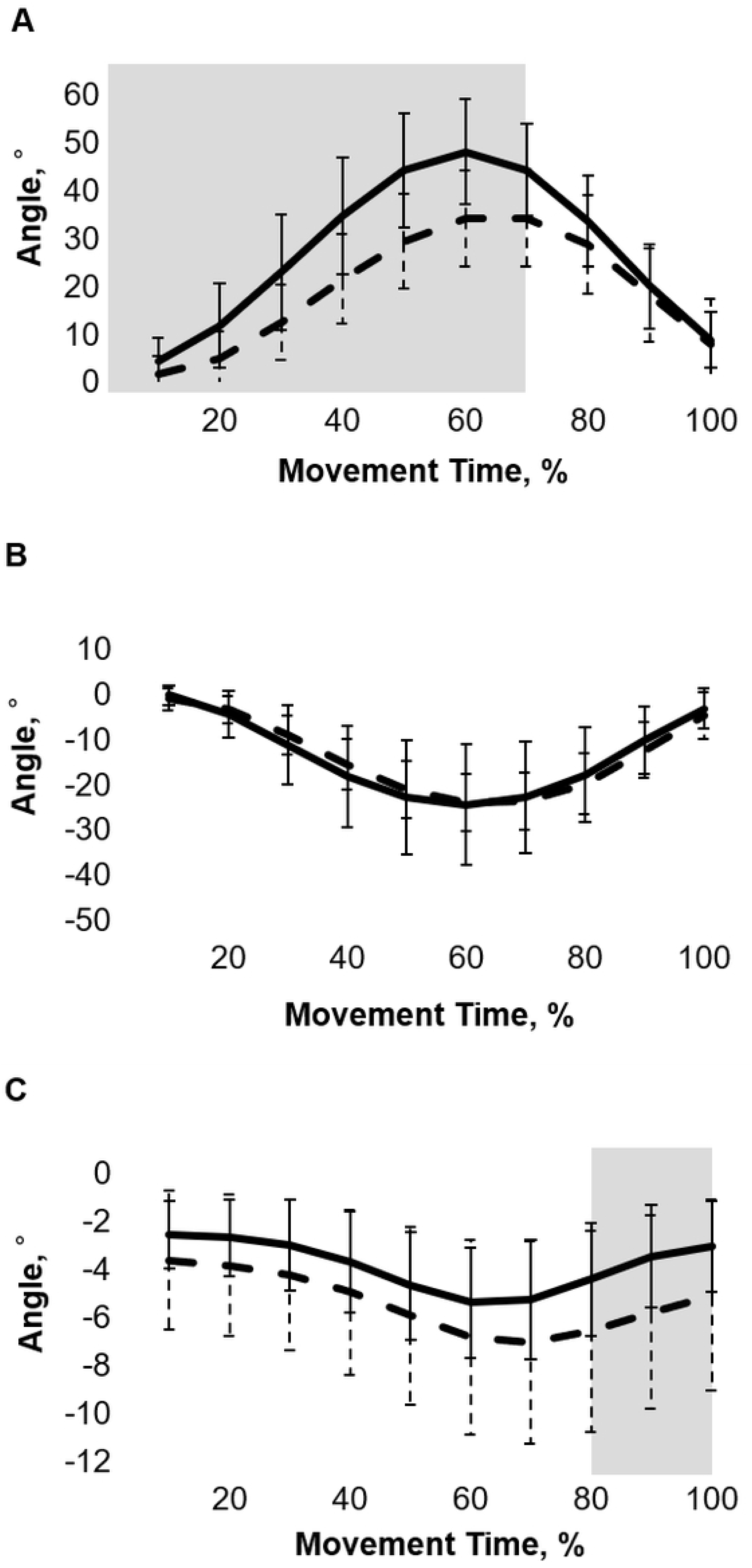
Lower limb angles during a half squat. Data are presented as mean ± standard deviation. A) Thigh angle, B) Shank angle, C) Foot angle. Gray box areas indicate significant group differences between the copers and controls (*P*< .05). Solid line: controls; dotted line: copers.

**Fig 2.**
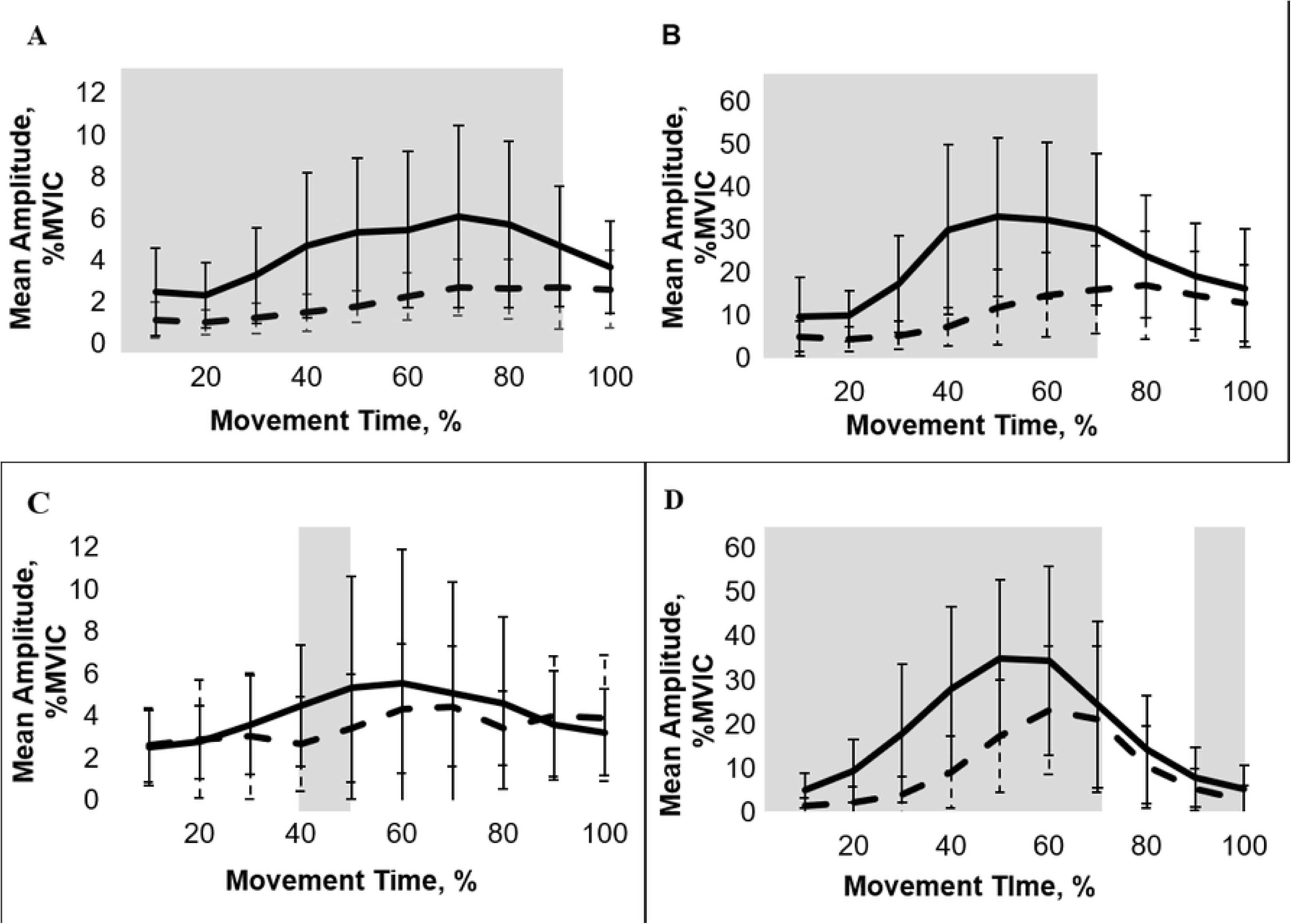
Muscle activities during a half squat. Data are presented as mean ± standard deviation. A) Biceps femoris long head, B) Rectus femoris, C) Gastrocnemius, D) Tibialis anterior. Gray box areas indicate significant group differences between the copers and controls *(P* < 0.05). Solid line: controls; dotted line: copers; MVIC: maximal voluntary isometric contraction. For gait, we observed no significant differences *(P* > 0.05) in lower limb angles and all four muscle activities between the copers and controls (Fig 3, Fig 4).

## Discussion

We examined lower limb rotational angles and muscle activities during half squats and gait between copers and healthy controls using IMUs and EMG. Our principal finding was that the copers had a reduced thigh rotational angle and less muscle activity during half squats, which supports our hypothesis. However, contrary to our hypothesis, there were no group differences in lower limb angles and muscle activities during gait.

The thigh angle was significantly reduced in copers in the half squat descending phase, whereas we observed no significant differences in the shank and foot angles during the same period (Fig 1). In other words, the copers had lesser knee bending during half squats than the control group.

However, there were no differences in the lower limb angles between the copers and controls during gait (Fig 3). Our finding that copers demonstrated less thigh angle during a half squat is inconsistent with previous findings that increased knee and hip angle were observed in people with CAI during movements that require foot fixation, such as stop jump or side-cutting movement [8, 13, 27]. After a first ankle sprain, the ligament and articular proprioceptors are damaged and lead to secondary nerve injury [3]. Since people with CAI reveal less ankle sagittal plane displacement due to damaged proprioceptors, they use compensatory strategies that increase knee and hip flexion to improve stability or to absorb the landing impact [13, 27, 28]. In addition, people with CAI may voluntarily limit their movements to avoid the positions in which they experience instability [13]. These compensatory strategies are more prominent in the group with great ankle laxity and less self-confidence in joint function [13, 28]. However, based on our results, copers may also restrict their thigh rotation unconsciously during half squat regardless of whether copers experience instability and if limb angles during gait have recovered to almost the same as that of controls without a history of ankle sprains.

**Fig 3.**
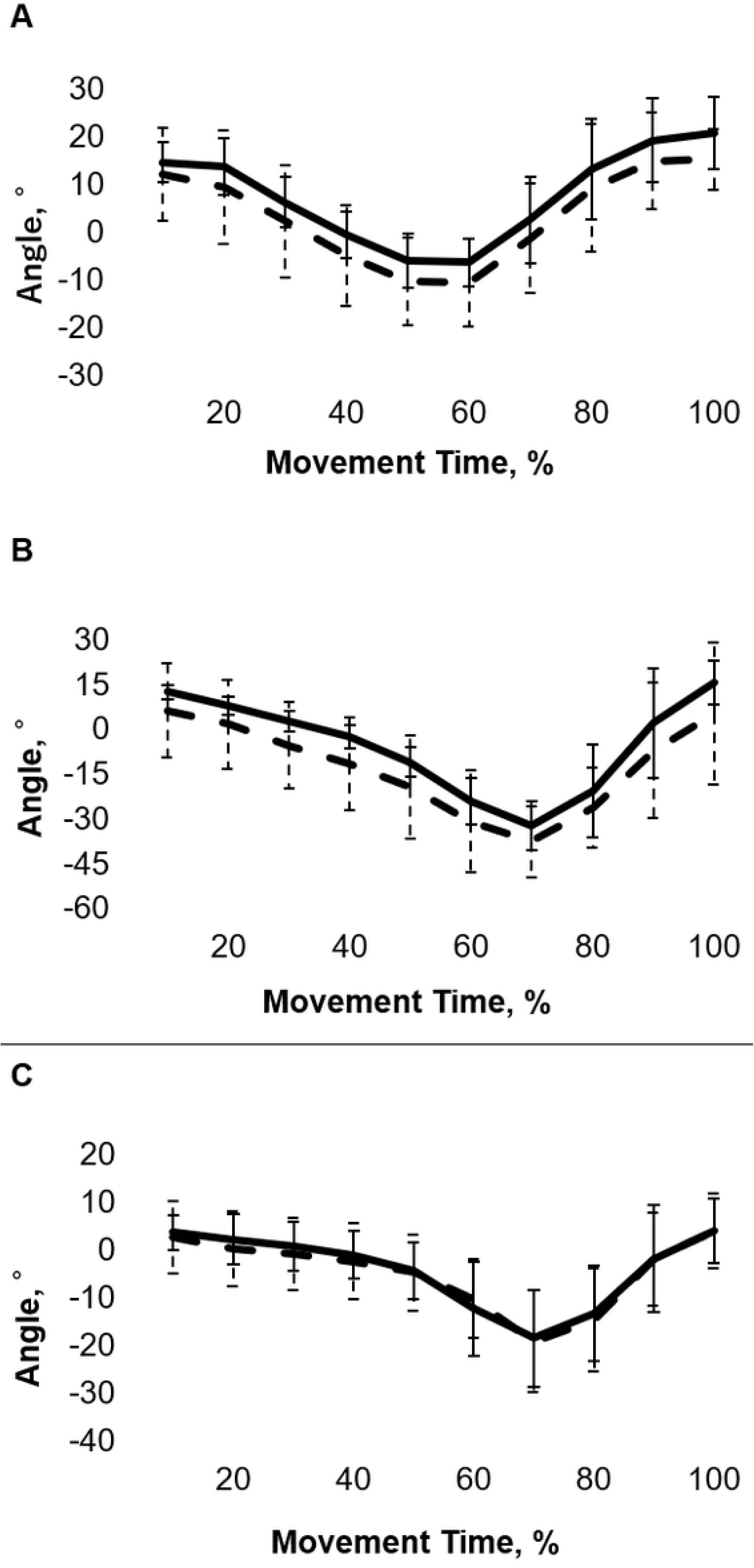
Lower limb angles during gait. Data are presented as mean ± standard deviation. A) Thigh angle, B) Shank angle, C) Foot angle. Solid line: controls; dotted line: copers.

The BFL, RF, and TA revealed large group differences between the copers and controls in the descending phase of the half squat (Fig 2). However, we observed no significant group differences in muscle activity during the gait (Fig 4). Less muscle activity during half squats is partially consistent with previous studies revealing that individuals with CAI exert lower muscle activity than controls during functional exercising tasks [1, 6, 7]. Copers showed lower activities of the RF and TA than controls during the same period of thigh restriction in half squats (Fig 1, Fig 2). TA activity is required to prevent the body from falling backward when bending the knee during a squat [19]. If the thigh angle is restricted due to TA activity weakness during weight-bearing movements, the dorsiflexion fixation of the foot at heel contact may be insufficient [29], resulting in a difference in foot angle during gait. However, there were no significant differences in the TA activity and foot angle during gait. Thus, these findings suggest that copers do not require TA activity compared to controls when involuntarily restricting thigh rotation during half-squats. As a result of unconscious thigh rotation restriction, the RF activity of the copers was also lower than that of the controls. The activity of RF, which is the target muscle of squatting exercise, plays an important role in squats when decreasing the gravitational acceleration and changing the movement direction from flexion to extension [19, 21, 22]. Therefore, copers may not obtain the desired exercise effect sufficiently during half squats compared to the effect obtained by healthy controls.

**Fig 4.**
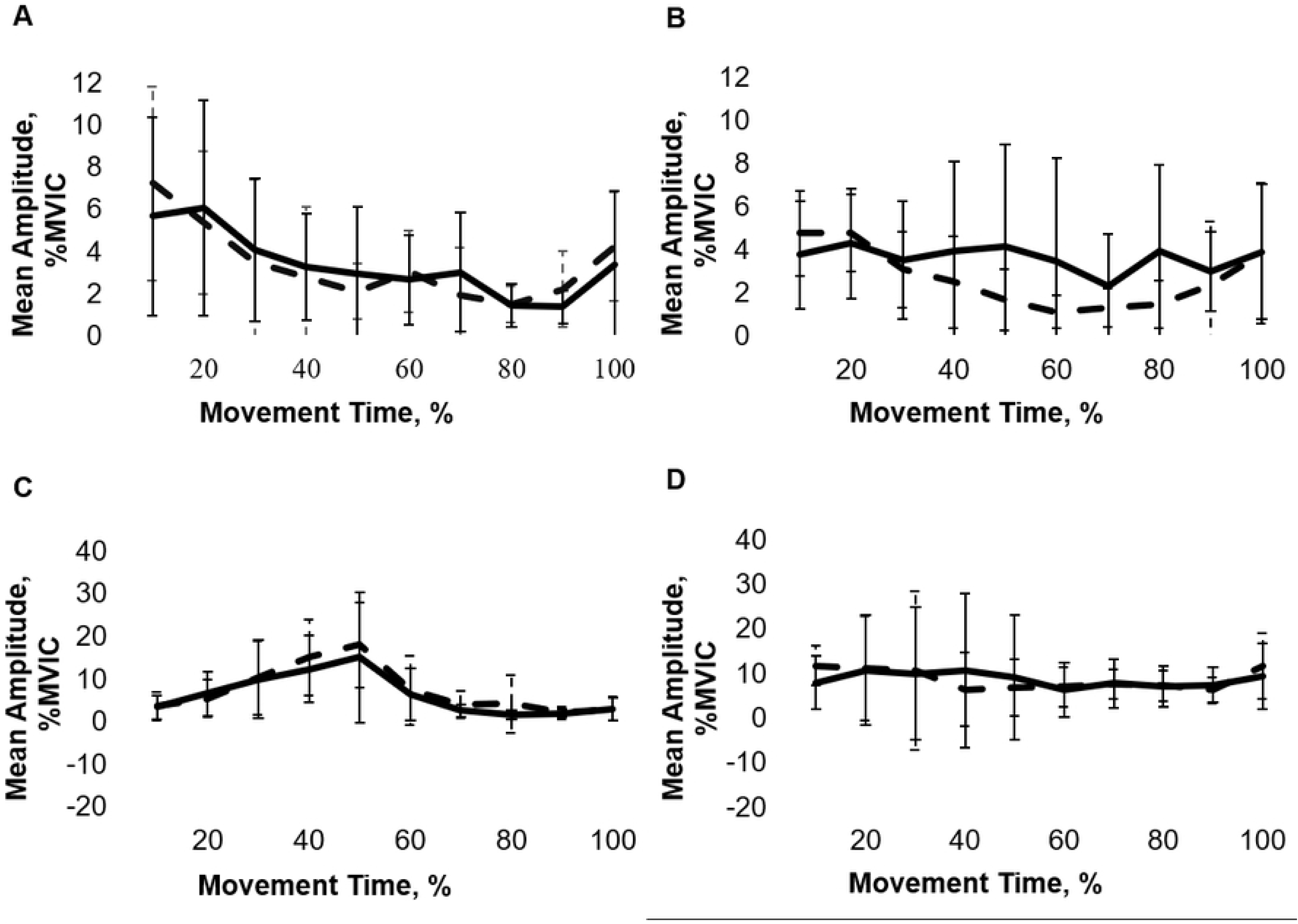
Muscle activities during gait. Data are presented as mean ± standard deviation. A) Biceps femoris long head, B) Rectus femoris, Gastrocnemius, D) Tibialis anterior. Solid line: controls; dotted line: copers; MVIC: maximal voluntary isometric contraction.

Our findings have several clinical advantages. When clinicians or coaches prescribe squatting exercises to copers, they need to determine whether copers are unintentionally restricting lower limb rotation. If copers unconsciously restrict lower limb rotation during squatting exercises, clinicians or coaches should remove the unconscious restriction using quantitative feedback from wearable sensors. Since the sensors used are small, portable, and cost-effective, they are suitable for use in clinical and sports settings. By removing the unconscious restriction of lower limb rotation, the targeted muscle activity of the intervention may increase, which may allow for more effective training. Thus, clinicians and coaches should focus on lower limb angles and muscle activities when prescribing exercise to people with a history of ankle sprain, even if they do not experience instability and there is no interference with their activities of daily living.

This study had some limitations. First, we were unable to recruit patients with CAI. Copers can adopt reorganization of the sensorimotor system by changing their movement strategies. In contrast, people with CAI fail to compensate for the loss of proprioception, which causes the recurrence of sprain and feelings of instability [10, 13]. To determine the cause of CAI development, future research is needed to identify the differences in kinematics and muscle activities among copers prospectively. Second, we examined only limb rotations in the sagittal plane and the four major muscle activities involved in knee flexion/extension and ankle dorsiflexion/plantarflexion. Great ankle frontal plane displacement and less activity of the peroneus longus are risk factors for recurrent sprains and progression to CAI [8, 9, 13, 27, 28]. Since copers showed angular displacements in the frontal plane compared to that in controls [8, 27], further research is needed to fully understand the features of movement strategies used by copers. Third, we examined only two simple movements: half-squats and gait. Verification with more advanced and complicated movements, such as running and jumping, should be considered for sports applications. Finally, we selected half squats, not full squats, to minimize head and trunk movements. A more comprehensive evaluation, including head and trunk movement evaluation, is desirable [18]. However, this was a basic study using wearable sensors to identify the movement strategies of copers; hence, a simplified model was used.

## Conclusions

Copers exhibited a restricted thigh rotational angle and less muscle activity compared with those exhibited by controls during half squats. There were no group differences in the lower limb angles and muscle activities during gait. Removing the unconscious restriction of the lower limb observed in copers with sensor-based quantitative feedback can provide therapeutic insights for copers to obtain more efficient training. Approximately 50% of copers return to their previous physical activity without appropriate treatment interventions. However, our study suggests the importance of assessing lower limb movements and muscle activities for copers to obtain efficient training effects, even if they do not experience instability and inconvenience in their daily lives.

## Acknowledgments

We would like to thank Editage (www.editage.com) for English language editing.

